# NanoPack2: Population scale evaluation of long-read sequencing data

**DOI:** 10.1101/2022.11.28.518232

**Authors:** Wouter De Coster, Rosa Rademakers

## Abstract

**Summary:** Increases in the cohort size in long-read sequencing projects necessitate more efficient software for quality assessment and processing of sequencing data from Oxford Nanopore Technologies and Pacific Biosciences. Here we describe novel tools for summarizing experiments, filtering datasets and visualizing phased alignments results, as well as updates to the NanoPack software suite.

**Availability and implementation:** Cramino, chopper, and phasius are written in Rust and available as executable binaries without requiring installation or managing dependencies. NanoPlot and NanoComp are written in Python3. Links to the separate tools and their documentation can be found at https://github.com/wdecoster/nanopack. All tools are compatible with Linux, Mac OS, and the MS Windows 10 Subsystem for Linux and are released under the MIT license. The repositories include test data, and the tools are continuously tested using GitHub Actions.

## Introduction

Long-read sequencing from Pacific Biosciences (PacBio) and Oxford Nanopore Technologies (ONT) has evolved from single genomes and small groups of individuals to large population-scale cohorts (Beyter *et al*., 2021; De Coster *et al*., 2021). Simultaneously, also the increasing economic cost and climate impact of computational tasks necessitate more efficient bioinformatic methods for data quality assessment and processing (Pereira *et al*., 2017). Several tools have been developed for the quality assessment of long-read sequencing data, however without scaling to populations of >100 genomes (Leger *et al*., 2020; Watson *et al*., 2015; Lanfear *et al*., 2019; De Coster *et al*., 2018). In this manuscript, we present newly developed tools that fulfill this need and efficiently assess characteristics specifically relevant to long-read genome sequencing, including alignments spanning structural variants and phasing read alignments. Phasing, i.e. assigning each sequenced fragment to a parental haplotype by identifying co-occurring variants (Martin *et al*., 2016; Edge and Bansal, 2019), is important in identifying potential functional variants in association studies and for the pathogenicity of putative compound heterozygous variation. Furthermore, we present an update on NanoPlot and NanoComp from the NanoPack tools (De Coster *et al*., 2018).

### Software description

Improvements to NanoPlot and NanoComp are, among several code optimizations, the generation of additional plots, using dynamic HTML plots from the Plotly library and as such enabling further exploration by the end users. The tools now also support using the programming-language agnostic Arrow data format as input. Chopper is a tool that combines the utility of NanoFilt and NanoLyse, for filtering reads based on quality, length, and contaminating sequences, and delivers a 7-fold speed up compared to the python implementation, making use of the Rust-Bio library (Köster, 2016) and Rust bindings to minimap2 (Li, 2018).

Summarization of long-read sequencing experiments using NanoStat (De Coster *et al*., 2018) is too slow considering the yields that are nowadays common with nanopore sequencing. Cramino, making use of rust-htslib (Bonfield *et al*., 2021; Köster, 2016), provides a much faster alternative for metrics based on the data output, mean coverage, the number of reads, their mean and median length, and sequence identity relative to the reference genome. Long reads span structural variants, and penalizing the read accuracy for a large gap is undesirable. For this reason, Cramino calculates the gap-compressed identity, defined as the edit distance relative to the read length, while counting consecutive gaps as just one difference. Cramino allows filtering on read length and optionally outputs a rudimentary evaluation of the karyotype and biological sex by calculating normalized read counts per chromosome, calculates the MD5 checksum to control for data integrity, and provides metrics of the read phasing performance. Importantly Cramino remains compatible with the rich visualizations from NanoPlot and NanoComp by generating output in the Arrow format, on top of the optional lightweight histograms for read length and read identity from Cramino itself. For a human genome with 50x coverage using 4 cores for BAM/CRAM decompression, Cramino takes 12 minutes with a peak memory usage of 147 Mb without optional output, or 21 minutes with a peak memory usage of 690 Mb for full output including histograms, karyotype, phasing metrics, MD5 checksum, and generation of the Arrow file.

Phasius is developed for the visualization of the results of read phasing, which shows in a dynamic genome browser style the length and interruptions between contiguously phased blocks from a large number of individuals together with genome annotation (Figure 1), for example, segmental duplications (Bailey *et al*., 2002). Phasius takes 26 seconds to generate the example figure for 92 individuals in a 10 megabase interval, with 8 parallel threads and a peak memory usage of 4.3Gbyte. For the example figure, reads were aligned with minimap2 (Li, 2018) and alignment phasing with longshot (Edge and Bansal, 2019).

**Figure 1:**
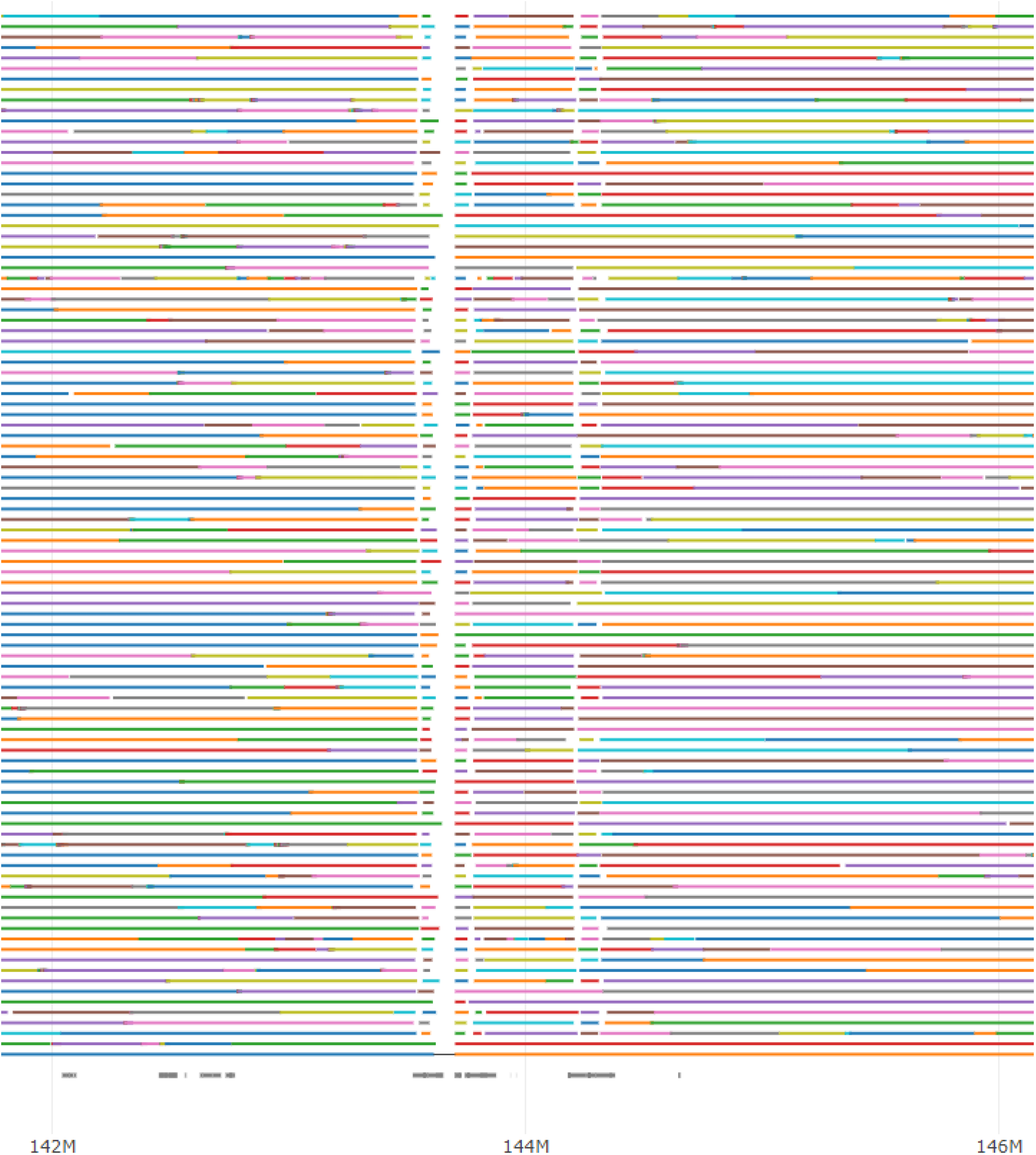
Example of Phasius output. This plot shows the haplotype phasing structure of chr7:142,000,000-146,000,000 for 92 individuals. Every horizontal line is from a single individual, with a change in color indicating the start of a new contiguously phased genomic segment. The annotation track (bottom) shows segmental duplications with grey bars, predictably breaking the phased blocks in the case of longer repetitive elements. An interactive example can be found at https://wdecoster.github.io/phasius

### Conclusion

NanoPack now offers tools ready for the evaluation of large populations with implementations in a more performant programming language, with a focus on features relevant to long-read sequencing. The software suite remains easy to install on all major operating systems and offers interactive visualization in HTML format.

## Acknowledgments

The authors acknowledge Ilias Bukraa for his contributions to NanoPlot and NanoComp.

